# Point mutations in topoisomerase I alter the mutation spectrum in *E. coli* and impact the emergence of drug resistance genotypes

**DOI:** 10.1101/621045

**Authors:** Amit Bachar, Elad Itzhaki, Shmuel Gleizer, Melina Shamshoom, Ron Milo, Niv Antonovsky

## Abstract

Identifying the molecular mechanisms that give rise to genetic variation is essential for the understanding of evolutionary processes. Previously, we have used adaptive laboratory evolution to enable biomass synthesis from CO_2_ in *E. coli*. Genetic analysis of adapted clones from two independently evolving populations revealed distinct enrichment for insertion and deletion mutational events. Here, we follow these observations to show that mutations in the gene encoding for DNA Topoisomerase 1 (*topA*) give rise to mutator phenotypes with characteristic mutational spectra. Using genetic assays and mutation accumulation lines, we show that point mutations in *topA* increase the rate of sequence deletion and duplication events. Interestingly, we observe that a single residue substitution (R168C) results in a high rate of head-to-tail (tandem) short sequence duplications, which are independent of existing sequence repeats. Finally, we show that the unique mutation spectrum of *topA* mutants enhances the emergence of antibiotic resistance in comparison to mismatch-repair (*mutS*) mutators, and lead to new resistance genotypes. Our findings highlight a potential link between the catalytic activity of topoisomerases and the fundamental question regarding the emergence of *de novo* tandem repeats, which are known modulators of bacterial evolution.

## Introduction

In clonal microorganisms, spontaneous mutations in the DNA sequence are a dominant source of genetic variation (Miller 1996). The rate and spectrum of mutational events are the outcome of extrinsic environmental conditions (e.g., exposure to DNA damaging agents) and intrinsic cellular processes (e.g., DNA replication fidelity). As natural selection acts on the genetic variation within a population, identifying mutagenic mechanisms is essential in understanding evolutionary outcomes, such as the emergence of antibiotic resistance (Chopra, O’Neill, and Miller 2003).

Population genetic models suggest that under strong selective conditions, such as exposure to novel environmental stress, mutator strains with high mutation rates can persist in the evolving population (Wielgoss et al. 2013; Good and Desai 2016). These mutators, which have spontaneous mutation rates of up to three times those of the ancestral strain, are frequently observed in laboratory evolution experiments (Sniegowski, Gerrish, and Lenski 1997; Bachmann et al. 2012; Phaneuf et al. 2019). A high prevalence of mutator strains is also found in natural bacterial populations, for example, in clinical isolates of *Pseudomonas aeruginosa* from the lungs of cystic fibrosis patients (Oliver et al. 2000), and among pathogenic isolates of *Escherichia coli* and *Salmonella enterica* (LeClerc et al. 1996).

The genetic basis of mutator strains is often traced to sequence variation that affects one of the cellular mechanisms related to DNA replication, repair, or maintenance. A partial list of mutator alleles includes the genes of the mismatch repair system, the oxidative lesion 8-oxodG system, and mutations affecting the proofreading capacity of DNA polymerases (Miller 1996). As the rate and spectrum of mutational events in mutator strains are shaped by the affected cellular mechanism (Foster et al. 2015), distinct mutational biases can affect the evolutionary dynamics of bacterial populations, as demonstrated in the case of antibiotic resistance (Couce, Guelfo, and Blázquez 2013).

## Results

### Biased mutation spectrum observed in two adaptive evolution experiments

We previously reported on the use of laboratory evolution in chemostats to adapt a metabolically engineered *E. coli* strain towards biomass synthesis from CO_2_ (Antonovsky et al. 2016; Herz et al. 2017). In two independent adaptive evolution experiments, we noticed a distinct bias in the types of mutation that were fixed in the population. Specifically, in clones isolated from the first evolution experiment we found that ≈85% of the detected mutations (28 of 33) were short sequence deletions, with deletion lengths of one to four base pairs (bps) (Supplementary Table 1). In a second independent experiment, sequencing of adapted clones revealed that over 60% of the acquired mutations (14 of 22) were base insertions. Notebly, all these insertion mutations were head-to-tail (tandem) sequence duplications (Supplementary Table 1). Tandem sequence duplication mutations are generally considered to arise from “slippage” replication events (Levinson and Gutman 1987; Zhou, Aertsen, and Michiels 2014), however, the newly formed duplicated sequences were independent of existing sequence repeats. A detailed description regarding the adaptive laboratory evolution process and the characterization of the evolved strains has been previously reported (Antonovsky et al. 2016; Herz et al. 2017).

To further explore the genetic basis underlying the biased mutation spectra that was observed in these two adaptive evolution experiments, we used whole-genome-sequencing to identify potential mutator genes in the adapted clones. We found that *topA,* encoding for DNA Topoisomerase 1 (TopA), was independently mutated in both of the evolving populations. In clones isolated from the first evolution experiment, where short sequence deletions dominated the mutation spectrum, we identified a point mutation in *topA* (g104c) that leads to a nonsynonymous substitution of an arginine residue (R35P). In the second evolution experiment, where a high frequency of tandem sequence duplication was observed, we found a point mutation (c502t) that results in the nonsynonymous substitution of another arginine residue (R168C). TopA belongs to the ubiquitous family of 1A topoisomerases, and is known to relieve helical torsion in the chromosome by introducing transient single-strand breaks in the DNA backbone. Previous studies highlight the involvement of topoisomerases in maintaining genome stability, for example by preventing the accumulation of DNA:RNA hybrid R-loops and the removal of DNA incorporated ribonucleotides (Cho, Kim, and Jinks-Robertson 2015; Usongo and Drolet 2014). Considering the key roles of topoisomerase in maintaining genome stability, we hypothesized that the point mutations identified in the evolving populations give rise to mutator phenotypes with a highly biased mutational spectrum.

### R35P and R168C Topoisomerase I mutants show biased mutation spectra, enriched in sequence deletions and insertions

To explore the potential effect of Topoisomerase 1 on the spectrum of mutational events, we genetically introduced the mutated *topA* alleles to an *E. coli* BW25113 genetic background (Methods). Whole-genome sequencing validated that the resulting strains, expressing either the R35P or the R168C mutated enzyme, were viable in the absence of compensatory mutations (Supplementary Table 2). However, we observed that mutations in *gyrA* or *gyrB* genes, with positive fitness effect, often emerged in the R35P strain during continuous strain propagation. Compensatory mutations in gyrase genes can reduce the accumulation of DNA torsional stress, and were reported to be essential for the viability of *topA* null mutants (Stockum, Lloyd, and Rudolph 2012).

Next, we quantified the spectrum of mutational events (i.e., point mutations, insertions and deletions) in the *topA* mutant strain s and in a *topA*^+^ strain using two independent drug resistance assays (Methods). We used 2-Deoxy-D-galactose (DOG), an inhibitor of galactokinase (encoded by *galK*), and azidothymidine (AZT), an inhibitor of thymidine/deoxyuridine kinase (encoded by *tdk*), as selection agents. Both drugs inhibit the growth of *E. coli* on minimal media via interactions with known molecular targets (Warming et al. 2005; Shelat, Parhi, and Ostermeier 2016). Briefly, we plated cells that were cultured in drug-free media on solid minimal media containing the toxic drug (either DOG or AZT) and a suitable carbon source (Methods). Resistant mutants that gave rise to colonies were isolated, and the resistance-conferring mutation in each colony was determined by sequencing of the known drug target gene. To ensure the analysis of independent mutational events, only a single colony was sampled from every assay repeat.

As shown in Fig. 1, sequencing of *galK,* the molecular target of DOG, revealed that resistant colonies arising from *topA* mutants exhibit significantly altered mutation spectrum in comparison to the *topA*^+^ strain (n=24 for each strain, Supplementary Table 3). In colonies arising from the R168C mutant strain, we found that 50% of the resistance conferring mutations in *galK* were sequence insertions, composed entirely from *de novo* tandem duplications with lengths of 2-19 bps. In the R35P mutant strain, we found that all of the resistance conferring mutations in *galK* were due to short sequence deletions, with lengths of 1-7 bps. These results were in marked contrast with the point mutation dominated spectrum (≈66%) that was observed in resistant colonies arising from the wild-type strain. When we repeated the experiment using AZT as a selection drug, the results were in agreement with the values reported above (Supplementary Figure 1 and Supplementary Table 3). In addition to the observed differences in the mutation spectra of *topA* mutants in comparison to the *topA^+^* strain, we noted that both R35P and R168C mutants exhibited strong localization pattern of mutations with well-defined hotspots (Fig. 1b). This contrasts with the uniform spatial distribution of mutations observed in the wild-type strain, where mutations occurred across the *galK* coding sequence without noticeable hotspots. Similarly, a mutational hotspot was also observed in the *topA* mutants when AZT was used a selection drug, where ≈50% of the resistance conferring mutations were localized in a 20 bps region (Supplementary Fig. 1).

**Figure 1.**
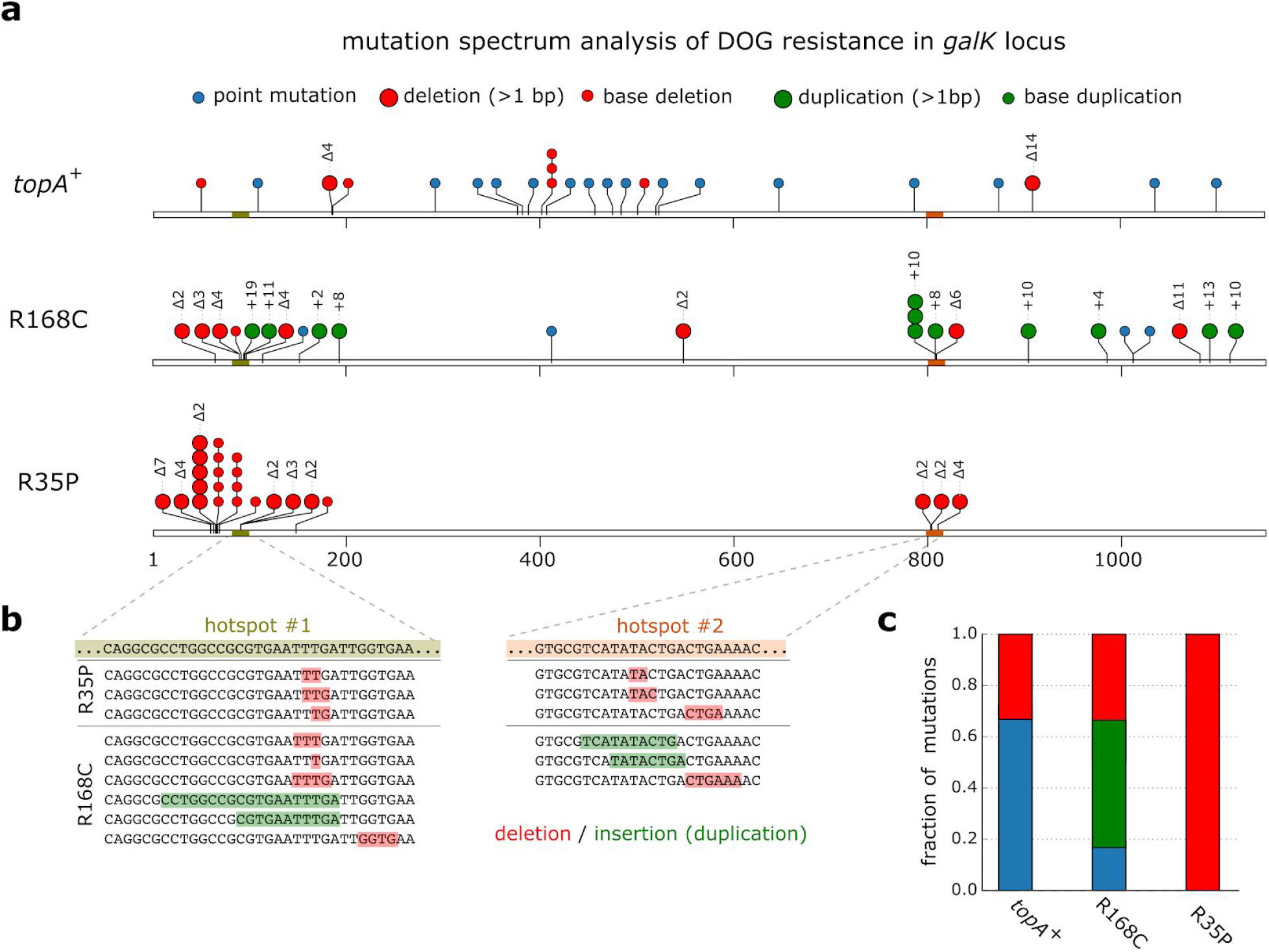
Drug resistance assay reveals an enrichment of insertion and deletion mutations in *topA* mutants. (a) Mutation spectrum analysis of DOG resistant colonies arising in R168C and R35P mutant strains or from *topA^+^* strain. To determine the resistance conferring mutations, we used Sanger sequencing of PCR amplicons of the *galK* locus. Overall, we sampled 24 independent resistant colonies for each genetic background. Notably, only a single colony was sampled from every assay to ensure that the analysis of independent mutational events. (b) Resistance conferring mutations arising in the *topA* mutants were mostly localized in two mutations hotspots in *galK.* (c) In contrast to the wild-type strain, where the majority of resistance conferring were due to point mutations, *topA* mutants are highly enriched in deletions (R35P) and tandem sequence duplications (R168C).

### Estimation of genomic mutation rates in R168C and R35P Topoisomerase I mutants using mutation accumulation lines

To quantify the genome-wide mutation rate of *topA* mutants, we conducted a mutation accumulation assay in which isogenic replicates were passaged through a single-colony bottleneck for 1200 or 600 generations. We used whole-genome sequencing to identify the mutations that accumulated in each line and calculated the rates of deletion, insertion, and point mutations for the different strains (Supplementary Table 4). As shown in Fig. 2, our analysis revealed that the rate of short sequence deletion events in the R35P mutant is significantly higher in comparison to the *topA*^+^ strain (p-value < 0.001, t-test) with an approximate 100-fold change in the mean rate of short deletions.

**Figure 2.**
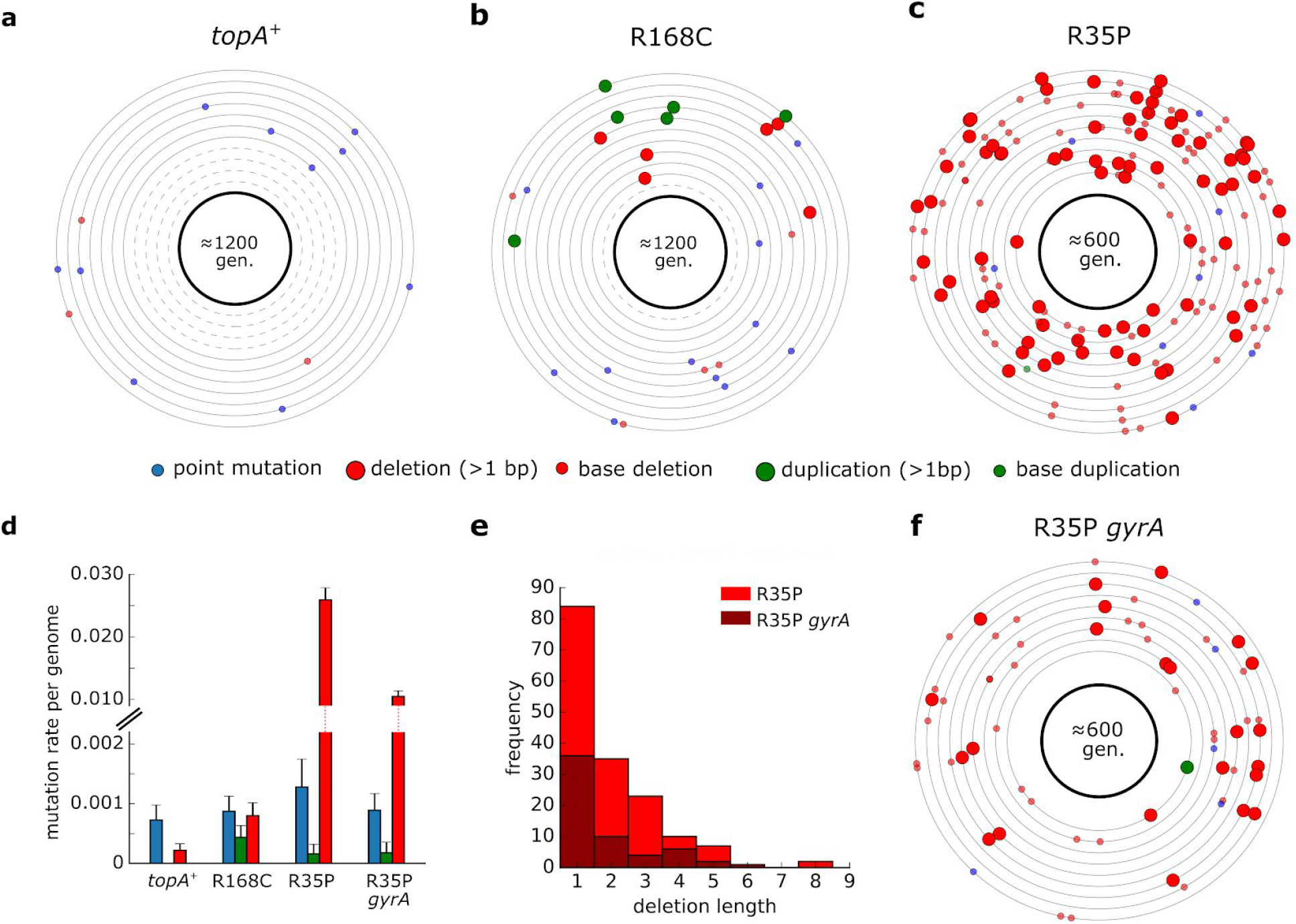
Rate and molecular spectrum of mutations in *topA* and wild-type strains as estimated by mutation accumulation lines. (a-c) Each circle represents the genome of a single mutation accumulation line (n=11). Mutations that were identified by whole-genome sequencing are positioned on the circle according to their chromosomal coordinates (clockwise). Dash circles indicate genome in which no mutation was detected. Detailed list of mutations can be found in Supplementary Table 4. (d) Average mutation rate per genome replication calculated for each mutation type. Error bars represent SEM. (e) Histogram of short deletion lengths that were identified in R35P mutant strains. (f) Mutations identified in isogenic lines (n=9) of R35P strain that contains a compensatory mutation in *gyrA.* Compensatory mutations in gyrase genes in *topA* mutants were reported to decrease the accumulation of DNA torsional stress and are essential for the viability of *topA* null mutants.

Since no sequence duplication events were observed in the *topA*^+^ lines during our experiment, we cannot directly determine the change in frequency of tandem duplication events. Previously published estimations of sequence insertion rate in wild-type *E. coli* found it to be ≈1×10^−4^ mutations per genome per generation (Lee et al. 2012), a rate which is 5 fold-lower than the rate we observed in the R168C mutant. However, this estimation of insertion rate in the wild-type strain is dominated by the rate of short sequence insertions (1-3 bps), occurring mainly in homopolymer tracts. As longer insertions, particularly those outside the context of existing sequence repeats, are significantly rarer, we note that the increase in the rate of *de novo* duplication mutations in the R168C is likely to be significantly higher. In contrast to the marked changes in the frequencies of deletion and duplication events, we found no significant difference in the rate of point mutations between the mutant and the *topA*^+^ strains. The point mutation rates measured for all of the strains in our analysis were in agreement with previously reported values of ä1-2×10^-3^ mutations per genome per generation (Lee et al. 2012). In addition, we used mutation accumulation lines to measure the effect of spontaneously arising compensatory mutations on the mutation rate in *topA* mutants. We isolated a clone in which a spontaneous mutation in *gyrA* had emerged in the R35P strain background. Based on our mutation accumulation analysis (n=9), we found that the rate of short sequence deletions in this double mutant remained significantly higher than the *topA*^+^ strain (p-value <0.001, t-test). However, the mutator phenotype was attenuated with an approximately ≈2 fold decrease in the deletions rate in comparison to the R35P lacking the compensatory mutation. Taken together, our results unequivocally show that R35P and R168C substitutions in TopA give rise to mutator phenotypes with distinct mutational bias.

### R35P and R168C substitutions affect highly conserved residues in DNA Topoisomerase I, lead to slower doubling time, and inhibit DNA relaxation activity

Sequence alignment of Topoisomerase 1 bacterial homologs shows that both R35P and R168C substitution mutations affect conserved residues in the protein (Fig. 3a). Based on crystal structures on the TopA, in was found that R168 participates in a network of ionic and hydrogen bond interactions which hold the DNA substrate in proper conformation for cleavage or re-ligation (Zhang, Cheng, and Tse-Dinh 2011). To determine the fitness effect of these mutations, we compared the growth rate of *topA* mutants to the native strain in LB and in glucose supplemented minimal media (Fig. 3b & Supplementary Figure 2). We found that in all cases, mutations in *topA* resulted in fitness cost (p-value <0.01 for all mutant strains, t-test). The effect was most prominent in the R35P mutant, where average doubling time was ≈50% longer in comparison to the *topA*^+^ strain. The compensating mutation in *gyrA* in the R35P background reduced the effect but did not eliminate the fitness cost. The smallest fitness cost was observed in the R168C mutant strain with ≈5% increase in doubling time in LB media. Next, we tested the effect of R35P and R168C mutations on the catalytic activity of the enzyme using *in vitro* DNA relaxation assay of heterologously expressed TopA mutant enzymes. For both of the mutant variants we observed a decrease in plasmid relaxation activity in comparison to the native enzyme. In the R168P mutant, we found that the minimal amount of enzyme required for complete plasmid relaxation was 4-fold higher in comparison to the wild-type enzyme, while in the R35P mutant protein we were not able to detect any plasmid relaxation activity.

**Figure 3.**
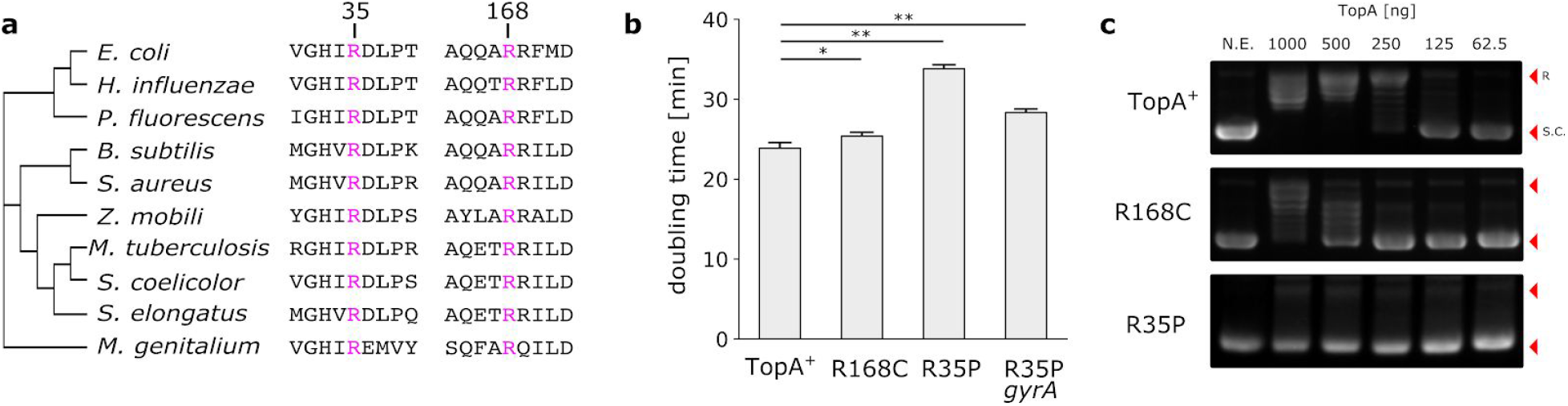
R168C and R35P mutations affect conserved residues and decrease supercoiling relaxation activity. (a) Multiple sequence alignment of bacterial topoisomerases indicating that the mutated residues (labeled in pink), R35 and R168, are highly conserved. (b) Doubling times of *topA^+^* stain in comparison to *topA* mutants in LB media. Error bars represent standard deviation of biological repeats (n=6). * p < 0.01, ** p < 0.001 (c) *In vitro* plasmid relaxation assay shows significant reduction in DNA supercoiling activity in R168C and R35P in comparison to the native enzyme.

### The biased mutations spectra in *topA* mutants impact the emergence of antibiotic resistance

Previous studies have shown that the mutational spectra in mismatch repair *E. coli* mutator strains affect the distributions of beneficial mutations in an antibiotic resistance model system (Couce, Guelfo, and Blázquez 2013). As the mutational spectrum defines accessible beneficial mutations, we sought to experimentally study the effect of the biased mutation spectra observed in *topA* mutants on the emergence of antibacterial resistance. We performed fluctuation assays to compare the rate of spontaneously arising resistants to D-Cycloserine (DCS), a broad-spectrum antibiotic, in the R35P mutant strain and in an MMR-deficient mutator strain (ΔmutS). Previous studies demonstrated that *mutS* mutators exhibit over 100-fold increase in the frequency of point mutations (Foster et al. 2018). This value is comparable to the rate of deletion mutations observed in the R35P strain, although the two strains differ in their mutation spectra (point mutations versus short sequence deletions). We find that under our experimental conditions, DCS resistant colonies emerged from ≈50% of R35P *topA* mutant cultures, but only from ≈3% of the *mutS* cultures (Fig. 4a, n=96). Whole-genome sequencing of independently arising resistant colonies from R35P strain cultures (n=5) revealed that all of the analyzed clones acquired mutations in *ispB,* an essential gene in the biosynthesis of isoprenoid quinones. Specifically, we observed the repeated occurrence of complex mutations, combining bases insertions and deletions, in a localized hotspot in the *ispB* gene. Sanger sequencing of *ispB* gene verified that in all five occurrences, these mutations impacted 2-3 amino acids within the coding sequence (Fig. 4b and Supplementary Table 5). In contrast, resistant colonies arising from cultures of the *mutS* strain (n=5) were dominated by point mutations with no single locus that was mutated in more than two clones (Supplementary Table 5). The marked difference in DCS resistance rates and genotypes between the *mutS* and *topA* mutators strengthen previous observations regarding the key role of the mutational spectra on the emergence of antibiotic resistant phenotypes.

**Figure 4.**
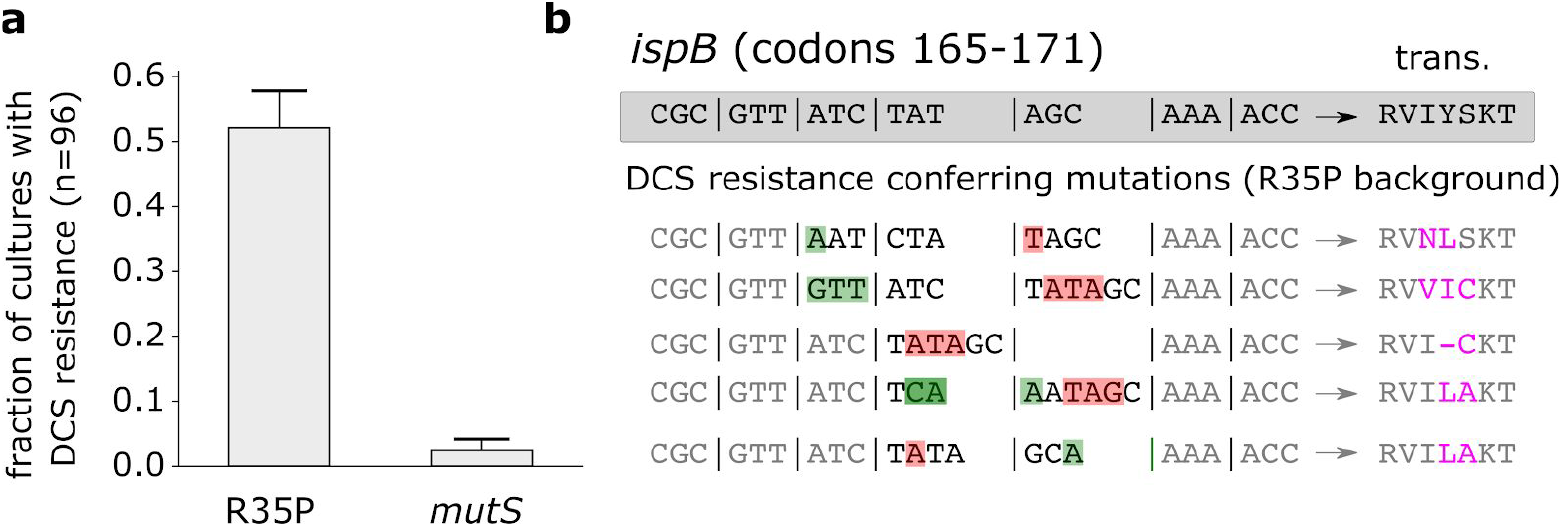
Cycloserine resistance conferring mutations in *ispB* observed in *topA* mutant (R35P) but not in MMR mutator *(mutS).* (a) Drug resistance fluctuation assay of R35P *topA* mutant and the *topA^+^* stains. Error bars represent the standard deviation of three experimental repeats, with 96 cultures samples per experiment. (b)Whole genome sequencing of cycloserine resistant colonies arising in the R35P *topA* mutant revealed complex mutations in *ispB* (n=5). In all five resistant clones, at least 2 residues in the protein sequence were affected by a combination of sequence deletions (red) and insertions (green).

## Discussion

Identifying the cellular processes that give rise to mutagenesis is crucial for the mechanistic understanding of genetic variation. Here, we show that mutations perturbing the relaxation activity of DNA topoisomerase I in *E. coli* lead to mutator phenotypes with deletion and tandem-duplication enriched mutation spectra. Potential sources of the mutagenesis observed in *topA* mutants include the direct involvement of the enzyme, for example by the formation of a stable cleavage complex, or through a secondary mechanism linked to the accumulation of torsional strain due to the impaired activity of mutants. Extensive studies in yeast associated short sequence deletions within tandem repeats with the mutagenic processing of ribonucleotides by Topoisomerase I (Kim et al. 2011; Huang, Ghosh, and Pommier 2015; Kim and Jinks-Robertson 2017). In *E. coli*, disruption of *topA* leads to excessive negative supercoiling and the accumulation of RNA:DNA hybrid R-loops and the stabilization of non B-DNA forms (Drolet 2006). As both of these non-canonical DNA structures are suggested to stimulate genetic instability, this highlights a potential mechanism for the mutator phenotype in *topA* strains. For example, a recent study demonstrated that dCas9-induced R-loop formation lead to the accumulation of deletion, insertion and complex mutations in yeast (Laughery et al. 2019). While our results unequivocally identify mutations in *topA* as the genetic basis of mutator phenotypes in *E. coli*, further studies are required to elucidate the molecular details underlying the difference in mutation rates and spectra between R35P and R168C *topA* mutants.

Fixation of *topA* mutants had been previously observed in multiple laboratory evolution experiments (Phaneuf et al. 2019). For example, in the long-term evolution experiment in *E. coli,* mutations in *topA* with positive fitness effects emerged in multiple evolving populations (Crozat et al. 2005). However, to the best of our knowledge, this is the first description of a *topA* mutator phenotype with an accelerated rate of tandem duplications in *E. coli.* Tandem sequence repeats are a well-documented source of genetic instability that play a significant role in microbial evolution (Treangen et al. 2009). While several molecular mechanisms pertaining to tandem repeat expansion and contraction have been described (Levinson and Gutman 1987; Bichara, Wagner, and Lambert 2006), the mechanism underlying *de novo* formation of tandem repeats from non-repeating sequences remains unknown. Our findings highlight a link between the catalytic activity of DNA topoisomerase I and the emergence of *de novo* tandem repeats, a ubiquitous source for genetic instability.

## Acknowledgments

The authors wish to thank Yinon Bar-On, Dr. Dan Davidi, Roee Ben Nissan, Prof. Rotem Sorek and Prof. Zvi Livneh for the helpful discussions. This research was supported by the Yeda – Sela Center for Basic Research, European Research Council (Project NOVCARBFIX 646827); the Israel Science Foundation (grant No. 740/16); Beck-Canadian Center for Alternative Energy Research; R.M. is the Charles and Louise Gartner professional chair.

## Methods

### Strains and Genomic Modifications

An *E. coli* BW25113 strain (Grenier et al., 2014) was used as the parental strain for all further genomic modifications. The evolved hemiautotrophic strains in which the R35P and R168C *topA* mutations were originally identified, have been previously described (Antonovsky. et al., 2016). For the construction of *topA* mutants (referred to in the text as strains R35P and R168C) in the background of the BW25113 strain we used P1 transduction according to the following procedure. First, P1 lysate was prepared from the ΔsohB780::kan Keio strain (JW1264-2, referred in the text as topA+ strain), in which a kanamycin resistance marker is located in proximity (≈1Kb) to the *topA* locus. The lysate was used to transduce the evolved hemiautotrophic strains containing *topA* mutations, and the transduced cells were plated on Kanamycin supplemented LB plates (25 ug/ml). Next, we used Sanger sequencing of the *topA* amplicons to screen for transduced clones in which the resistance marker has been introduced into the *sohB* locus without affecting the mutation in the adjacent *topA* gene. Positive clones, in which the resistance cassette and mutated *topA* locus have been genetically linked, were used as donor strains for a second round of P1 transduction into a wild-type BW25113 strain. The short distance between the resistance marker and the *topA* locus (<1Kb) in comparison to the large size (~100kb) of the DNA fragment delivered to the recipient facilitated the allelic exchange of the mutated *topA* locus to the recipient BW25113 strain. Whole-genome-sequencing was used to validate the introduction of the mutated *topA* genes, along with the ΔsohB780::kan marker, to an otherwise unperturbed genetic background in the BW25113 strain. The ΔsohB780::kan Keio strain (JW1264-2), containing the native *topA* sequence, was used as a control strain in all of the genetic assays (referred in the text as *topA*^+^ strain). *sohB* is a non-essential gene that encodes polypeptide S49 peptidase family protein. No phenotypes related to mutation rate and spectrum are associated with the ΔsohB780::kan genotype. The strain referred to in the text as ΔmutS is the ΔmutS738::kan from the KEIO collection (JW2703-2).

### Drug resistance assays

All assays were performed in 96-well microplates. A single colony from each strain was inoculated into M9 minimal media supplemented with glycerol (2 g/l) as carbon source and suitable antibiotics (25 ug/ml kanamycin). Following overnight incubation (37°C, 220 RPM) the saturated culture was diluted to a concentration of ≈10^5^ cells/ml based on OD measurements, and 10 μL (≈10^3^ cells) were inoculated into 150 μL of M9 minimal media supplemented with limiting amount of glycerol (40 mg/L) to control the overall number of cell divisions. The culture was incubated for 24-48 hours, after which 5 μL drops from each well were plated on M9 agar plate supplemented with glycerol (2 g/L) and the counter-selection drug. The assay was repeated twice, using either 2-Deoxy-D-galactose (DOG) at a final concentration of 2 mg/mL (Sigma-Aldrich, D4407) or Azidothymidine (AZT) at a final concentration of 5 μM (1.3 μg/mL) (Sigma-Aldrich, A2169) as the counter-selection drugs. The agar plates were incubated for 48 hours and resistant colonies were sampled to determine the resistance conferring mutation in each colony. Notably, only a single colony was sampled from each culture to ensure the analysis of independent mutational events. For each sampled colony, we used PCR to amplify and Sanger sequence the known genes (galK/DOG and *tdk/AZT*) whose loss-of-function render resistance to the drug. The resulting sequences were compared to the wild-type sequence of *galK* or *tdk* genes and mutations in the resistant colonies were annotated using Geneious software (http://www.geneious.com). Assays for DCS resistance were performed similarly to the previously described, with the following modifications: (a) LB media was used (b) DCS concentration was 133 mg/L (c) Mutation calling was performed by whole-genome-sequencing of resistant colonies as described below.

### Mutation accumulation lines

For each strain, a glycerol stock was inoculated in LB media, incubated overnight, and streaked on an LB agar plate supplemented with 25 μg/mL kanamycin. Isogenic lines were established from single, well isolated colony. Lines were propagated daily by streaking a random single colony from each line on LB agar plates supplemented with 25 μg/mL kanamycin. The lines were maintained for 48 passages (≈1200 generations) for *topA*^+^ and R168C strains or 24 passages (≈600 generations) for strains R35P and R35P gyrA. As the growth rates of the different strains varied, different incubation times were needed to obtain comparable generations per passage rates, as determined by CFU counting of typical colonies (≈25 generations/passage). Freezer stocks were sampled from the founding strains and every 25 passages. Whole-genome sequencing was used to determine the mutations accumulated in the final clones, as detailed below. The mutation rate per genome replication was calculated for each mutation type based on the average number of mutations across all lines for each strain.

### Whole-genome-sequencing and mutation calling

A single colony was inoculated in 3mL of LB media with kanamycin (25 ug/mL) and incubated to saturation. The culture was harvested by centrifugation and genomic DNA was extracted using DNeasy Blood and Tissue Kit (QIAGEN). Sequencing libraries were prepared using Nextera tagmentation reaction as described by Baym et al., 2015. The final libraries were sequenced using HiSeq 2500 or NextSeq 500 sequencers (Illumina) to yield single reads with a typical coverage of x30 per sample. Demultiplex reads were mapped to the reference of E. coli BW25113 (CP009273.1) and mutations were called using breseq pipeline version 0.25 (Deatherage and Barrick, 2014). We note that our analysis does not capture certain types of mutations, such as copy number variations.

### Purification of recombinantly expressed TopA variants

His-tagged TopA proteins (R35P, R168C and wild-type) were expressed in *E. coli* BW25113 transformed with the TopA encoding plasmid pCAN24-6xHis-topA obtained from the ASKA library (Kitagawa et al., 2005). The R35P and R168C encoding mutations were introduced to the coding sequence of topA using PCR based site-directed-mutagenesis according to standard protocols. The resulting plasmids, pCAN24-6xHis-topA:R35P and pCAN24-6xHis-topA:R168C, were sequenced to validate the mutated sequence. For recombinant protein expression, a single colony of E. coli BW25113 cells, transformed either with the native or one of the mutated plasmids, was inoculated in 3 mL of LB media supplemented with chloramphenicol (34 μg/mL). Following overnight incubation (37 °C, 220 RPM) the saturated culture was diluted 1:20 into 50 mL of fresh media. TopA over-expression was induced at mid-exponential phase (≈0.5 OD) by the addition of IPTG (0.5mM). The cells were harvested and the protein was purified using His-Spin Protein Miniprep kit (Zymo Research) according to the manufacturer’s instructions. Following protein purification, we used Amicon Ultra 30K centrifugal filter (Millipore) to replace the elution buffer with the plasmid relaxation assay reaction buffer. Protein purity was validated using polyacrylamide gel electrophoresis and GelCode Blue staining (Thermo Fisher). Purified protein concentration was determined using the bicinchoninic acid (BCA) assay (Thermo Fisher) according to manufacturer’s protocol.

### Plasmid relaxation assay

To assay for one unit of relaxation activity, purified enzymes of the same concentrations were diluted serially, ranging from 1000–62.5 ng and assayed for DNA relaxation activity in a final reaction volume of 10 μL with 40 mM Tris-HCl (pH 8.0), 50 mM KCl, 10 mM MgCl_2_, 100 μg/mL BSA, and 800 ng of supercoiled plasmid DNA. The reaction was incubated at 37°C for 30 min, after which it was terminated by the addition of SDS to a final concentration of 1%. The DNA was electrophoresed in a 0.8% (w/v) agarose gel with TAE buffer for 45 minutes at 135V. The gel was stained with ethidium bromide, washed twice in DDW for 1 hour, and photographed under UV light. We estimated the relative activity of according to the defined least quantity of the enzyme required for complete relaxation of negatively supercoiled DNA under the given reaction conditions.

